# A protein-trap allele reveals roles for *Drosophila* ATF4 in photoreceptor degeneration, oocyte maturation and wing development

**DOI:** 10.1101/2021.05.18.444691

**Authors:** Deepika Vasudevan, Hidetaka Katow, Grace Tang, Hyung Don Ryoo

## Abstract

Metazoans have evolved various stress response mechanisms to cope with cellular stress inflicted by external and physiological conditions. The Integrated Stress Response (ISR) is an evolutionarily conserved pathway that mediates adaptation to cellular stress via the transcription factor, ATF4. Loss of function of *Drosophila* ATF4, encoded by the gene *cryptocephal* (*crc*), results in lethality during pupal development. The roles of *crc* in *Drosophila* disease models and adult tissue homeostasis thus remain poorly understood. Here, we report that a protein-trap MiMIC insertion in the *crc* locus generates a crc-GFP fusion protein that allows visualization of crc activity in vivo, and acts as a hypomorphic mutant that uncovers previously unknown roles for *crc*. Specifically, the *crc* protein-trap line shows crc-GFP induction in a *Drosophila* model for Retinitis Pigmentosa (RP). This *crc* allele renders photoreceptors more vulnerable to age-dependent retinal degeneration. *crc* mutant adult animals also show greater susceptibility to amino acid deprivation and reduced levels of known crc transcriptional targets. Furthermore, this mutant allele shows defects in wing veins and oocyte maturation, uncovering previously unknown roles for *crc* in the development of these tissues. Together, our data establish physiological and pathological functions of *crc-*mediated ISR in adult *Drosophila* tissues.

## Introduction

Virtually all organisms have evolved stress response mechanisms to mitigate the impact of homeostatic imbalance. The Integrated Stress Response (ISR) pathway, conserved from yeast to humans, is one such mechanism initiated by stress-responsive eIF2α kinases. ISR pathway has been linked to the etiology of a number of human diseases including neurodegenerative disorders, diabetes, and atherosclerosis, amongst others (Chan et al. 2016; Ivanova and Orekhov 2016; Back et al. 2012; Ma et al. 2013). There is thus significant interest in better understanding the ISR signaling pathway.

Each ISR kinase responds to a different type of stress: PERK, an ER-resident kinase, responds to disruption in endoplasmic reticulum (ER) homeostasis (*e*.*g*. misfolding proteins, calcium flux); GCN2, a cytoplasmic kinase, responds to amino acid deprivation; PKR, a cytoplasmic kinase, responds to double stranded RNA, and; HRI, a cytoplasmic kinase, that responds to oxidative stress (Donnelly et al. 2013). When activated by the corresponding cellular stress, the ISR kinases phosphorylate the same downstream target: the α-subunit of the initiator methionyl-tRNA (Met-tRNA_i_^Met^) carrying complex, eIF2. Such phosphorylation of eIF2α kinases leads to decreased availability in Met-tRNA_i_^Met^ resulting in lowered cellular translation (Sonenberg and Hinnebusch 2009). However, the translation of some mRNAs with unusual 5’ leader arrangements, such as the one encoding the ISR transcription factor ATF4, is induced even under such inhibitory conditions (Hinnebusch et al. 2016). ATF4 is a bZIP (basic Leucine Zipper) transcription factor that induces the expression of stress response genes, including those involved in protein folding chaperones, amino acid transporters, antioxidant genes (Back et al. 2009; Han et al. 2013; Fusakio et al. 2016; Shan et al. 2016).

The number of ISR kinases varies depending on organismal complexity, *e*.*g*. GCN2 in yeast, GCN2 and PERK in *Caenorhabditis elegans* (worms) and *Drosophila melanogaster* (flies), and all four ISR kinases in *Danio rerio* (zebrafish) and other higher vertebrates (Ryoo 2015; Mitra and Ryoo 2019). ATF4 remains the best-characterized transcription factor that is induced downstream of these kinases (Donnelly et al. 2013), and *Drosophila* has a functionally conserved ortholog referred to as *cryptocephal* (*crc*) (Fristrom 1965; Hewes et al. 2000). In addition to its well-characterized roles during cellular stress, a plethora of studies have demonstrated roles for ISR signaling components during organismal development (Pakos-Zebrucka et al. 2016; Mitra and Ryoo 2019). In *Drosophila*, both *Gcn2* and *Perk* mutants survive to adulthood (Kang et al. 2017; Vasudevan et al. 2020), and the emerging adults show phenotypes in the gut, wings and female ovaries (Wang et al., 2015; Armstrong et al. 2014; Malzer et al. 2018). On the other hand, *crc* mutants fail to reach adulthood (Fristrom 1965; Hewes et al. 2000). The *crc* hypomorphic point mutant, *crc*^*1*^, which causes a single amino acid change, results in delayed larval development and subsequent pupal lethality (Fristrom 1965; Hewes et al. 2000; Vasudevan et al. 2020). The most striking phenotype of the *crc*^*1*^ mutants is the failure to evert the adult head during pupariation, along with a failure to elongate their wings and legs (Fristrom 1965; Hewes et al. 2000; Vasudevan et al. 2020; Hewes et al. 2000; Gauthier et al. 2012).

The larval and pupal lethality of known *crc* alleles have however limited our understanding of *crc*’s roles in adult tissues. *crc* is cytogenetically close to the widely used FRT40 element, which has impeded efforts to study this mutation using FRT-mediated mitotic clones. Here, we report that a GFP protein-trap reporter allele in the *crc* locus acts as a hypomorphic mutant that survives to adulthood. We use this allele to discover that loss of *crc* results in accelerated retinal degeneration in a *Drosophila* model of autosomal dominant retinitis pigmentosa (adRP), a human disease whose etiology is linked to ER stress. Adult *crc* mutants show increased susceptibility to amino acid deprivation, consistent with what was previously known for GCN2. Additionally, we observe several developmental defects in adult tissues, including reduced female fertility due to a block in oogenesis. We also observe wing vein defects and overall reduced wing size in both male and female *crc* mutants.

## Results

### crc^GFSTF^ is a faithful reporter for endogenous crc levels

In seeking endogenous reporters of crc activity, we examined a “protein trap” line for *crc* generated as part of the Gene Disruption Project (Nagarkar-Jaiswal, DeLuca, et al. 2015; Nagarkar-Jaiswal, Lee, et al. 2015; Venken et al. 2011). The protein trap line is based on a MiMIC (<u>Mi</u>nos <u>M</u>ediated Integration <u>C</u>assette) element inserted randomly into various regions in the *Drosophila* genome. The cassette can be subsequently replaced with an EGFP-FlAsH-StrepII-TEV-3xFlag (GFSTF) multi-tag cassette using recombination mediated cassette exchange. One such insertion recovered through this project is in the intronic region of the *Drosophila crc* locus, which has been subsequently replaced with an EGFP-FlAsH-StrepII-TEV-3xFlag (GFSTF) multi-tag cassette using recombination mediated cassette exchange (Fig. 1). The splice donor and acceptor sequences flanking the cassette ensure that the GFSTF multi-tag is incorporated in the coding sequence of most abundantly expressed *crc* splice isoform, *crc-RA* (Hewes et al. 2000), to generate a multi-tag crc fusion protein (Fig. 1). Henceforth, this *crc* reporter allele is referred to as *crc*^*GFSTF*^, with the encoded fusion protein referred to as crc-GFP.

**Fig. 1.**
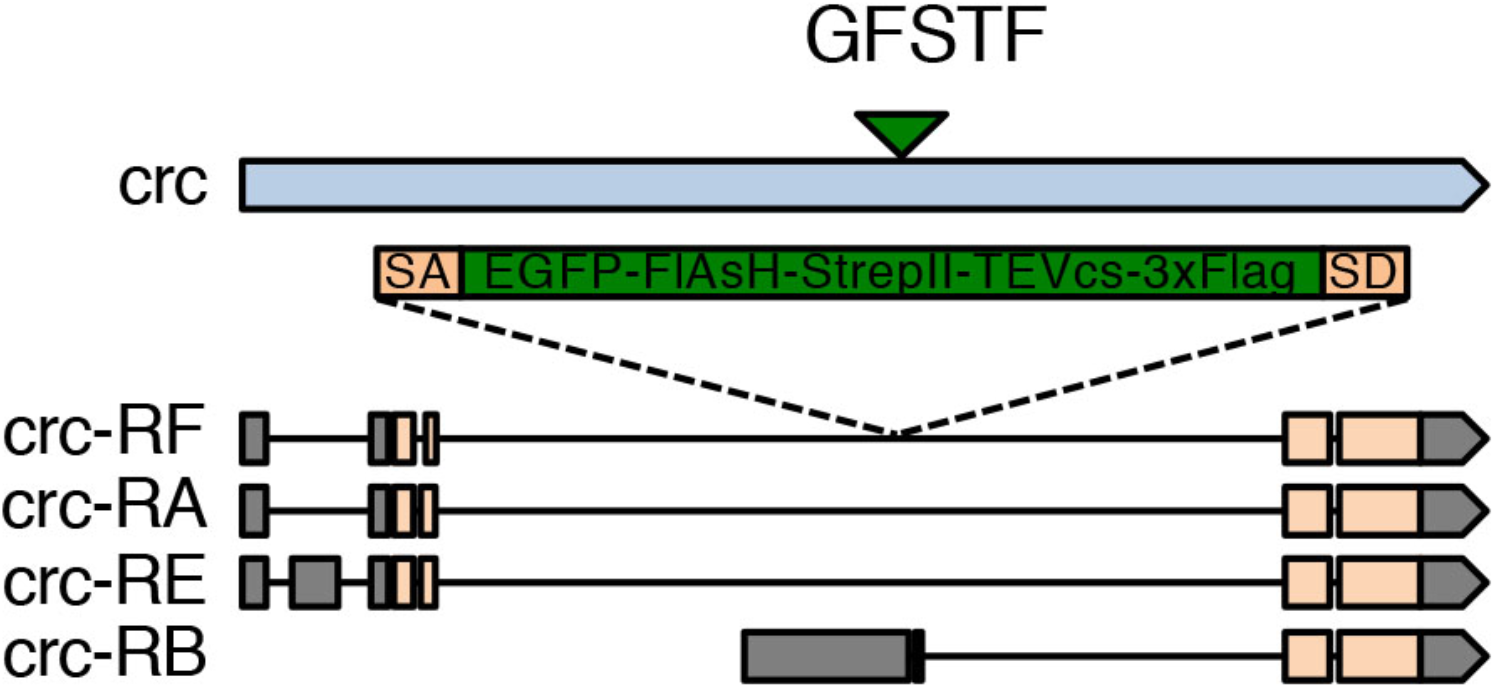
Schematic of the *crc* cytogenetic locus. The *crc* gene (blue bar) is known to encode at least four splice variants *crc-RA, -RB, -RE*, and *–RF*. These splice isoforms vary in their 5’ leader sequences (gray bars) and their coding exons (beige bars). MiMIC-mediated insertion of the GFSTF cassette in the genomic locus (green triangle) with a splice acceptor (SA) and splice donor (SD) sequences predicts the inclusion of a multi-tag exon (green box) in all the *crc* isoforms except *crc-RB*.

Our lab and others have utilized acute misexpression of Rh1^G69D^, an ER stress-imposing mutant protein, in third instar larval eye disc tissues using a *GMR-Gal4* driver (*GMR>Rh1*^*G69D*^) as a facile method to activate the *Perk-crc* pathway (Ryoo et al. 2007; Kang et al. 2015; Kang et al. 2017). We tested the utility of *crc*^*GFSTF*^ allele as an endogenous reporter for crc levels, and found robust induction of crc-GFP in third instar larval eye discs specifically in response to misexpression of Rh1^G69D^ protein but not control lacZ protein in the *crc*^*GFSTF*^/+ background (Fig. 2a, b). To validate that such induction was downstream of PERK activation by misfolding Rh1^G69D^, we generated *Perk* mutant FRT clones negatively marked by DsRed expression in the GMR compartment using ey-FLP. While control clones showed no change in the induction of crc-GFP (Fig. 2c), *Perk*^*e01744*^ mutant clones showed a complete loss of crc-GFP in *GMR>Rh1*^*G69D*^ eye imaginal discs (Fig. 2d). We also validated these observations in whole animal *Perk*^*e01744*^ mutants, where we observed a complete loss of crc-GFP in *GMR>Rh1*^*G69D*^ eye imaginal discs (Fig. S1a, b).

**Fig. 2.**
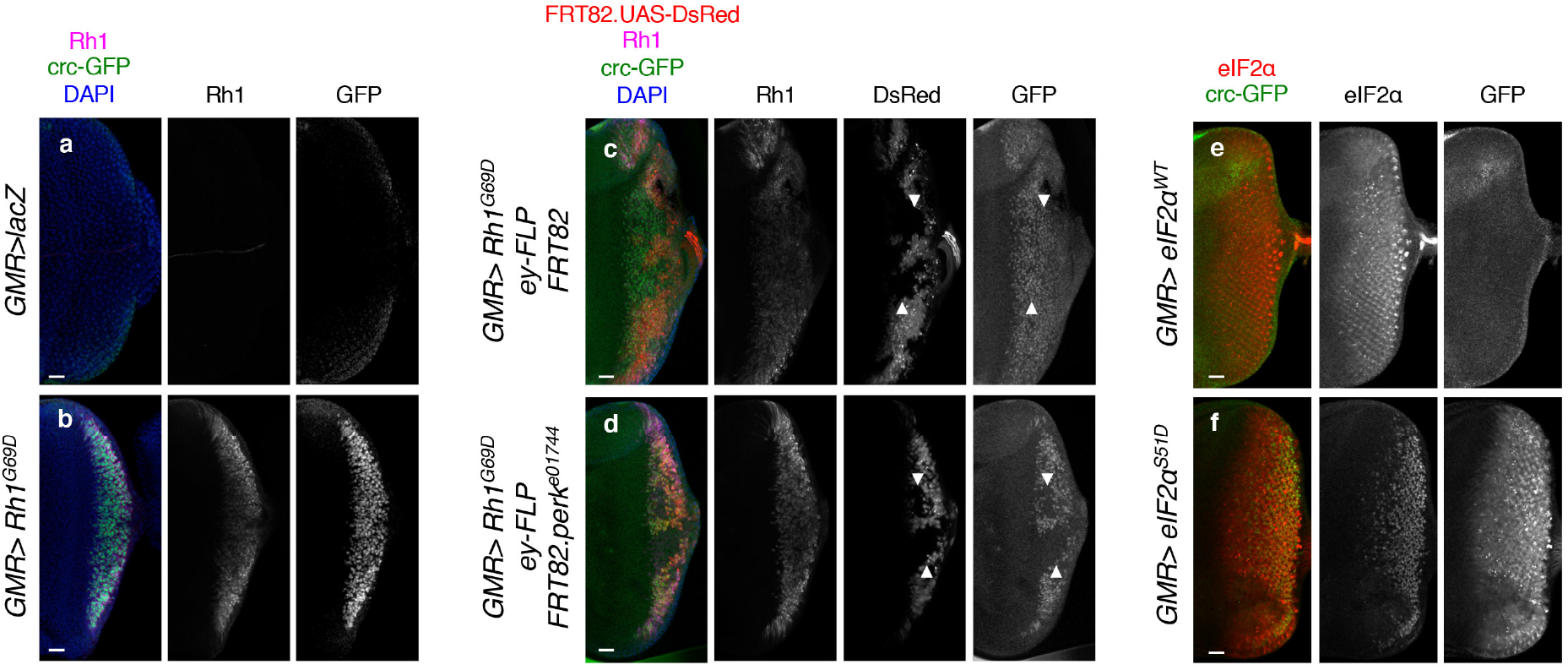
*crc*^*GFSTF*^ is a reporter for crc activity in vivo. a-b. Confocal images from eye imaginal discs from wandering third instar larva where *GMR-Gal4* drives the expression of either a control protein (*GMR>lacZ*) or mutant Rh1 (*GMR>Rh1*^*G69D*^), in the *crc*^*GFSTF*^*/*+ background. Here and in following images, the crc-GFP fusion protein was detected with anti-GFP (green), Rh1 was detected with 4C5 antibody (magenta), DAPI (blue) stains the nucleus. Rh1 and GFP single channels are shown separately in black and white; anterior is the left, posterior to the right; scale bars represent 25 μM. c-d. Confocal images of eye imaginal discs misexpressing Rh1^G69D^ (*GMR>Rh1*^*G69D*^) showing control clones (c, FRT82) and *Perk* mutant clones (d, *FRT82*.*perk*^*e01744*^) generated by eyeless-flippase (*ey-FLP*) in the *crc*^*GFSTF*^*/*+ background. Clones are negatively marked with DsRed (red). This DsRed expression was also driven by *GMR>* (white arrowheads). Rh1, DsRed and GFP single channels are separately shown in black and white images. The absence of GFP in DsRed negative clones demonstrate the effect of loss of *Perk* on crc-GFP induction in response to Rh1^G69D^. e-f. Confocal images of eye imaginal discs showing crc-GFP expression in response to wildtype eIF2α (*eIF2α*^*WT*^) or phospho-mimetic eIF2α (*eIF2α*^*S51D*^) ectopically expressed with *GMR-Gal4* (*GMR>*). Ectopic expression was confirmed by staining with anti-eIF2α (red). eIF2α and GFP single channels are shown in black and white.

Since the induction of crc in response to PERK activation occurs due to eIF2α phosphorylation (Sonenberg and Hinnebusch 2009), we examined whether crc-GFP induction we observed in Fig. 2a-d similarly occurs through this mechanism. Specifically, we generated a phospho-mimetic transgenic line where the Ser51 in eIF2α is mutated to Asp51 (*UAS-eIF2α*^*S51D*^). We also generated a corresponding control transgenic line containing wild type eIF2α (*UAS-eIF2α*^*WT*^). We next expressed these transgenes in flies containing *crc*^*GFSTF*^. While *GMR>eIF2α*^*WT*^ discs showed no detectable levels of crc-GFP, we found that *GMR>eIF2α*^*S51D*^ led to robust induction of crc-GFP in eye discs as detected by immunostaining with anti-GFP (Fig. 2e, f). These data demonstrate the applicability of *crc*^*GFSTF*^ as a reliable reporter of endogenous crc expression downstream of ISR activation.

### crc^GFSTF^ is a hypomorphic crc mutant allele

Similar to the previously characterized crc hypomorphic mutant allele, *crc*^*1*^, we observed that flies homozygous for *crc*^*GFSTF*^ exhibited a delay in head eversion and showed anterior defects (Fristrom 1965; Hewes et al. 2000). To further assess the effects of the *crc*^*GFSTF*^ allele, we performed a lethal phase analysis of development starting at the first instar larval stage. We found that a little over 50% of *crc*^*GFSTF*^ homozygotes were larval lethal (Fig. 3a), which is remarkably similar to larval lethality we previously reported for *crc*^*1*^ (Vasudevan et al. 2020). However, unlike *crc*^*1*^ homozygotes, only a small percentage of *crc*^*GFSTF*^ homozygotes showed prepupal and pupal lethality, with ∼30% of animals eclosing as adults (Fig. 3a). To ensure that these developmental defects cannot be attributed to background mutations in the *crc*^*GFSTF*^, we performed lethal phase analysis on *crc*^*GFSTF*^ in transheterozygotic combination with the hypomorphic *crc*^*1*^ allele. We found that *crc*^*GFSTF*^*/crc*^*1*^ transheterozygotes showed similar levels of larval and pupal lethality to *crc*^*GFSTF*^ homozygotes, with ∼25% of animals surviving to adulthood (Fig. 3a). These data together suggested that the *crc*^*GFSTF*^ allele may function as a *crc* loss-of-function allele.

**Fig. 3.**
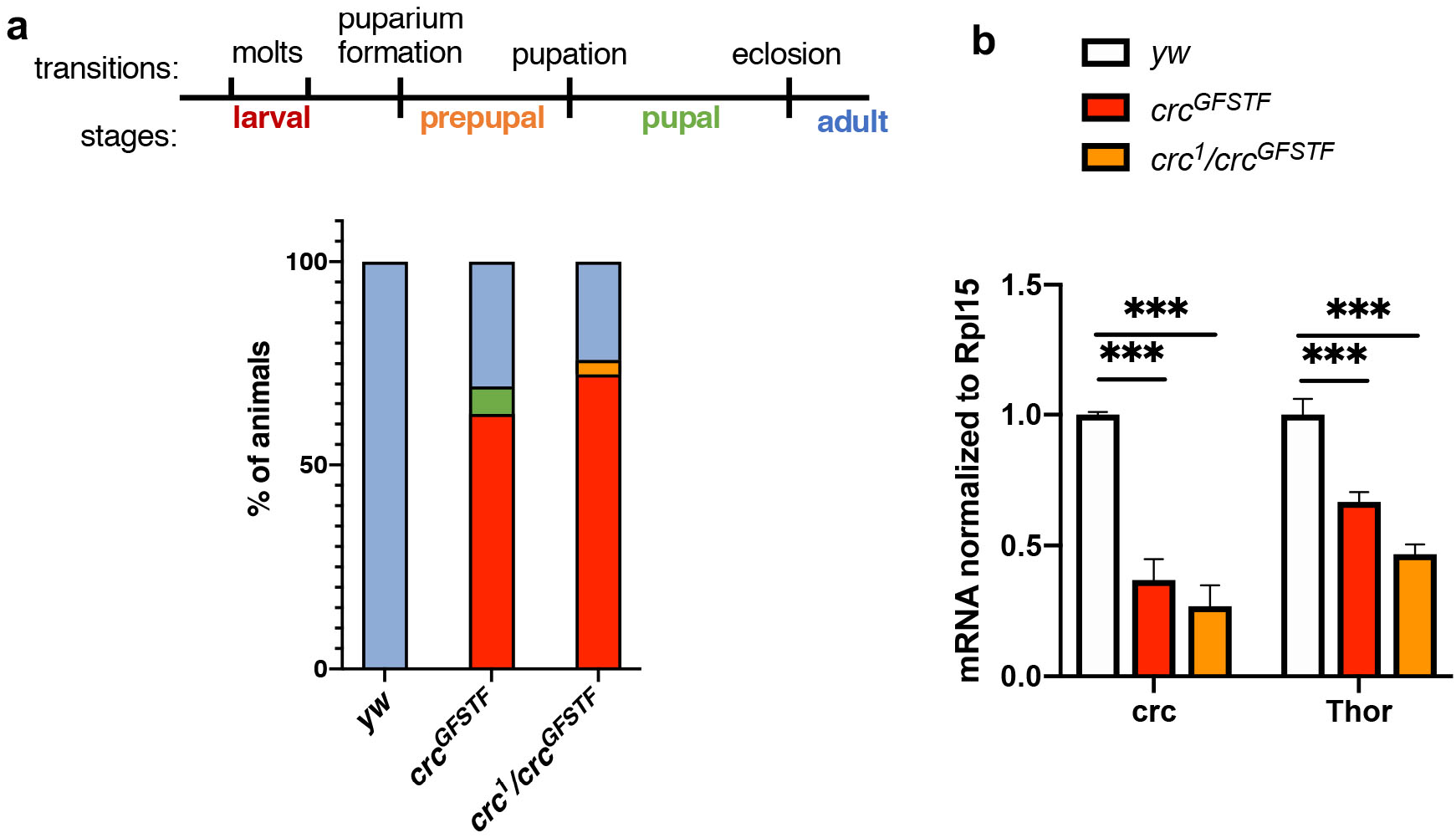
*crc*^*GFSTF*^ is a *crc* hypomorphic allele. a. (Top) Schematic showing transitions and stages during *Drosophila* development. (Bottom) Lethal phase analysis for control (*yw*), *crc*^*GFSTF*^ homozygotes, and transheterozygotes (*crc*^*GFSTF*^*/crc*^*1*^), color-coded per the schematic. n=100 for each genotype. b. qPCR analysis of *crc* and its transcriptional target, *Thor*, normalized to *Rpl15* from wandering 3^rd^ instar larval stages when crc expression is known to be elevated. Data represent the mean of 3 independent experiments, error bars represent standard error. ***= p<0.0001 calculated by a paired two-tailed *t*-test.

To examine if crc transcript levels are affected in *crc*^*GFSTF*^ mutants, we performed qPCR in the wandering 3^rd^ instar larval stage when crc activity is known to be high in fat tissues (Kang et al. 2015; Kang et al. 2017). We found that *crc*^*GFSTF*^ homozygotes show ∼65% decrease in *crc* transcript levels in comparison to control animals (Fig. 3b). We also tested crc activity by measuring mRNA levels of the well-characterized crc transcriptional target, *4E-BP* (*Drosophila Thor)*. We observed ∼40% lower levels of *Thor* in *crc*^*GFSTF*^ in comparison to control animals (Fig. 3b). This reduction in transcript levels of crc targets was also reproducible *crc*^*GFSTF*^/*crc*^*1*^ transheterozygotes (Fig. 3b). Taken together, these data indicate that *crc*^*GFSTF*^ acts as a mild hypomorphic mutant allele of *crc*.

### crc has a protective role in age-related retinal degeneration and amino acid deprivation

Nearly 30% of all adRP mutations are found in the Rhodopsin gene (Kaushal and Khorana 1994; Illing et al. 2002). Several of these Rhodopsin mutations impose stress in the ER (Kroeger et al. 2019). However, the role of ATF4 in adRP has remained unclear, and we sought to resolve this using the *crc*^*GFSTF*^ allele in a *Drosophila* model of adRP.

Clinically, adRP is characterized by age-related loss of peripheral vision, resulting in ‘tunnel vision’, and night blindness due to degeneration of rod photoreceptors (Kaushal and Khorana 1994). The *Drosophila* genome encodes several Rhodopsin genes, including *ninaE* that encodes the Rhodopsin-1 (Rh1) protein. The *ninaE*^*G69D*^ mutation captures essential features of adRP etiology: Flies bearing one copy of the dominant *ninaE*^*G69D*^ allele exhibit the age-related retinal degeneration as seen by photoreceptor cell death (Colley et al. 1995; Kurada and O’Tousa 1995). We found that *crc*^*GFSTF*^/*crc*^*1*^; *ninaE*^*G69D*^*/+* animals exhibited rapid retinal degeneration in comparison to *crc*^*GFSTF*^/*+; ninaE*^*G69D*^*/+* control animals, as monitored by pseudopupil structures in live flies over a time course of 30 days (Fig. 4a). While the earliest time point when control animals exhibit retinal degeneration is typically 13-15 days, *crc* homozygous mutant animals exhibited retinal degeneration as early as 2 days, with all animals displaying loss of pseudopupil structures by day 14 (Fig. 4a). Interestingly, we also found that *crc*^*GFSTF*^/*crc*^*1*^ animals exhibited age-dependent retinal degeneration even in the absence of *ninaE*^*G69D*^, indicating a protective role for crc in photoreceptors under physiological conditions during aging (Fig. 4a).

**Fig. 4.**
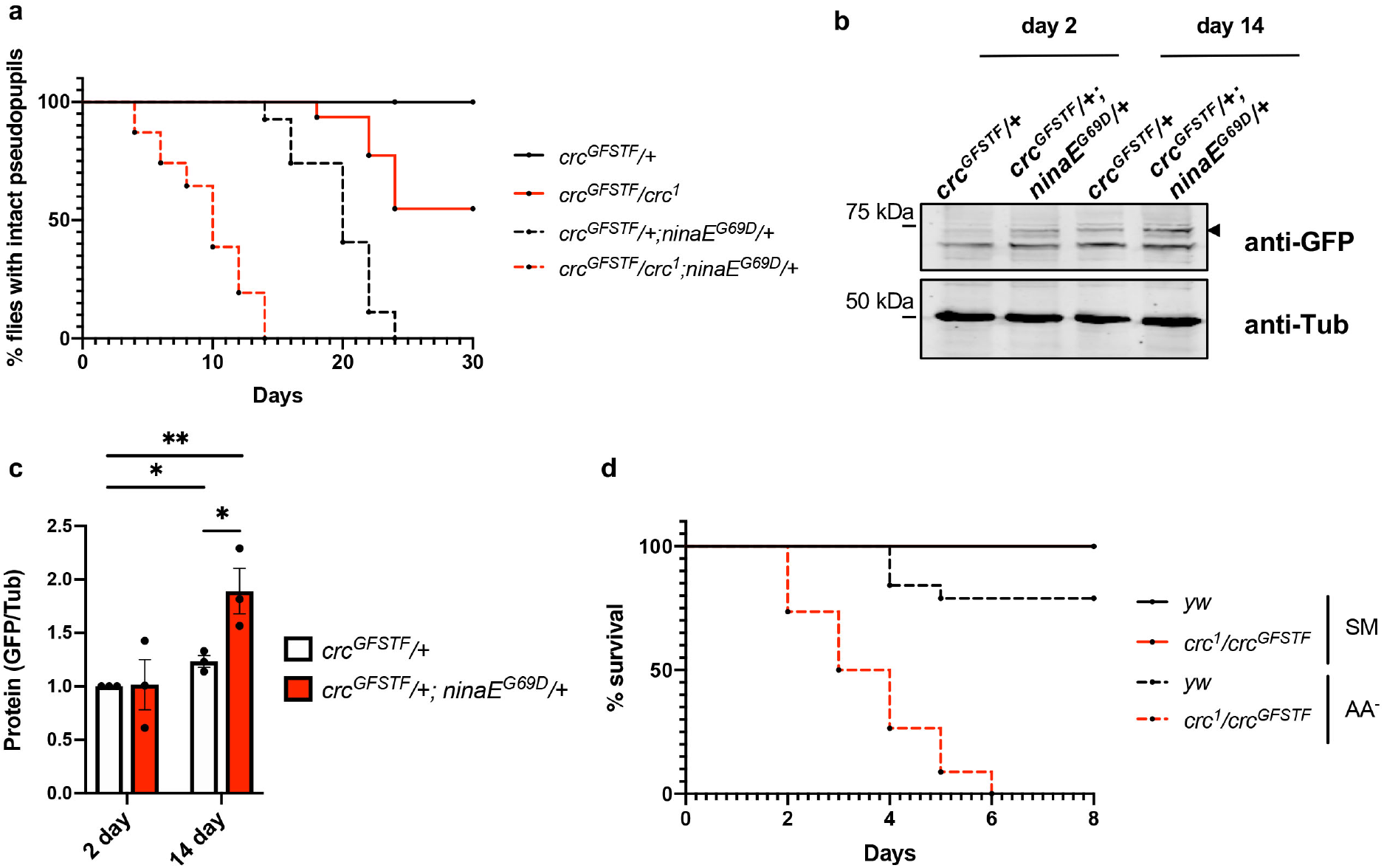
crc mediates *Perk* and *Gcn2* phenotypes in adult animals. a. Time course of pseudopupil degeneration in control and *ninaE*^*G69D*^/+ flies in *crc* heterozygote (*crc*^*GFSTF*^/+) and transheterozygous mutants (*crc*^*GFSTF*^*/crc*^*1*^). The difference in the course of retinal degeneration between the following pairs is statistically significant as assessed by the Log-rank (Mantel-Cox) test (p < 0.001): *crc*^*GFSTF*^/+ and *crc*^*GFSTF*^*/+*;*ninaE*^*G69D*^*/+, crc*^*GFSTF*^/*crc*^*1*^ and *crc*^*GFSTF*^*/crc*^*1*^;*ninaE*^*G69D*^*/+*, and, *crc*^*GFSTF*^/+ and *crc*^*GFSTF*^*/crc*^*1*^. (n = 100). b. Western blot analysis of fly head extracts from young (1-2 day) and aged (14-16 day) old flies of the control and *ninaE*^*G69D*^/+ animals also heterozygous for *crc*^*GFSTF*^. The upper panel shows the blot probed with anti-GFP to detect the crc-GFP fusion protein (distinguished by the black arrowhead) and lower panel shows Tubulin (anti-Tub) as a loading control. c. Quantification of western blotting data in (b) showing crc-GFP normalized to Tubulin. Data represent the mean from three independent experiments, error bars represent standard error. **= p<0.001, *=p<0.01 calculated by a paired two-tailed *t*-test. d. Time course of survival rate of adult females of indicated genotypes when fed with standard media (SM, solid lines) or amino acid deprived media (AA^-^, broken lines). Note that the curves for the flies fed SM for *yw* (solid black) and *crc*^*GFSTF*^*/crc* (solid red) overlap entirely. The difference in the survival rates between the following pairs is statistically significant as assessed by the Log-rank (Mantel-Cox) test (p < 0.001): *yw* (SM) and *yw* (AA^-^), *crc*^*GFSTF*^*/crc* (SM) and *crc*^*GFSTF*^*/crc* (AA^-^), *yw* (AA^-^) and *crc*^*GFSTF*^*/crc* (AA^-^). (n = 100).

To measure the expression of crc in aging photoreceptors, we performed western blotting of adult fly heads from young and old (2-week) flies to detect crc-GFP. While young control flies (*crc*^*GFSTF*^/+) showed very low levels of crc-GFP, flies bearing one copy of *ninaE*^*G69D*^ showed a substantial induction of crc-GFP (Fig. 4b, c). We observed that crc-GFP increases with age in both 2-week old control flies (*crc*^*GFSTF*^/+), with a concomitant increase in crc-GFP in *ninaE*^*G69D*^*/+* flies as well (Fig. 4b, c). These data substantiate the engagement of crc in photoreceptors in response to ER stress inflicted by the ER stress-imposing Rh1^G69D^, thus providing a basis for the protective roles of *Perk* in retinal degeneration.

In addition to rendering a protective effect during ER stress inflicted by Rh1^G69D^, we also tested if crc had an effect during amino acid deprivation in adult animals. We tested this by subjecting *crc*^*GFSTF*^/*crc*^*1*^ animals to amino acid deprivation by rearing animals on 5% sucrose-agar. While a majority of control animals survived up to 8 days, *crc*^*GFSTF*^/*crc*^*1*^ animals steadily succumbed to amino acid deprivation starting at day 2 with no survivors by day 6 (Fig. 4d). This is consistent with the idea that crc mediates the GCN2 response to amino acid deprivation in adult *Drosophila*.

### crc mutants show wing size and vein defects

*crc*^*GFSTF*^ provided an opportunity to examine previously unreported roles for *crc* in adult flies. We first observed that wings from both *crc*^*GFSTF*^ homozygotes and *crc*^*GFSTF*^/*crc*^*1*^ transheterozygotes showed a range of venation defects (Fig. 5a-c). The *Drosophila* wing has five longitudinal veins (annotated L1-L5) and two cross veins, anterior and posterior, ACV and PCV respectively (Fig. 5a). Severe wing defects in *crc*^*GFSTF*^ homozygous female and male flies were characterized by ectopic venation on L2, between L3 and L4, on L5, and also ectopic cross veins adjacent to the PCV (Fig. 5b, b’). *crc*^*GFSTF*^/*crc*^*1*^ transheterozygotes largely showed milder wing defects, characterized by ectopic venation on the PCV and on L5 (Fig. 5c, c’). We quantified these wing phenotypes in over forty animals of each sex and found that the penetrance and severity of the phenotypes were much stronger in females than in males (Fig. 5d). We also observed that *crc* mutant wings were smaller than in control animals (Fig. 5a-c). Quantification of wing area from animals of each sex revealed a statistically significant decrease in wing blade size in *crc*^*GFSTF*^ and *crc*^*GFSTF*^/*crc*^*1*^ males and females (Fig. 5e). To exclude the possibility of dominant negative effects of *crc*^*GFSTF*^, we also tested wings from *crc*^*GFSTF*^/+ heterozygotes but found no wing defects in these animals (Fig. S2). It is notable that *Gcn2* depletion in the wing reportedly causes venation defects (Malzer et al. 2018). Thus, our results suggest that *Gcn2*-mediated crc activation is involved in proper wing vein development.

**Fig. 5.**
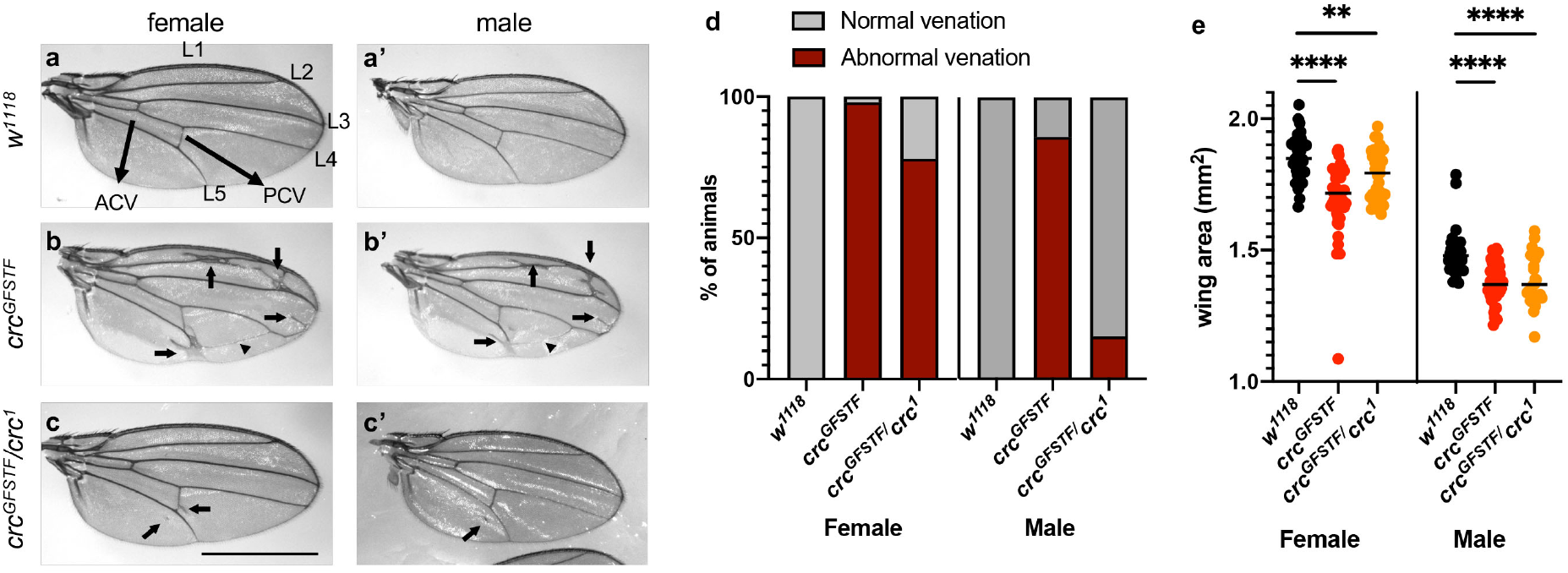
Adult *crc* mutants display developmental defects in the wing. a-c. Grayscale images of the right wing from female (a-c) or male (a’-c’) flies from the indicated genotypes. (a) shows the arrangement of wing veins in control (*w*^*1118*^) flies with L1-L5 marking longitudinal veins, and arrows marking the anterior cross vein (ACV) and posterior cross vein (ACV). Ectopic longitudinal veins in *crc*^*GFSTF*^ homozygotes and *crc*^*GFSTF*^*/crc*^*1*^ transheterozygotes (b, b’, c, c’) are marked by arrows and arrow heads point to ectopic cross veins. Scale bar= 1000μm. d. The penetrance of the ectopic vein phenotype in (a-c) quantified from 40 animals of each sex of the indicated genotypes. e. Area of the right wing from male and female flies of the indicated genotypes as measured in ImageJ. n≥27 for each genotype. Data represent the mean and error bars represent standard error. **= p<0.001, ****= p<0.00001 calculated by an unpaired two-tailed *t*-test.

### crc mutants exhibit decreased fertility due to defects in oogenesis

In trying to establish a stock of *crc*^*GFSTF*^, we observed that when mated to each other *crc*^*GFSTF*^ homozygotic males and females produced no viable progeny with very few of the eggs laid hatched to first instar larvae. To determine if this loss of fertility in *crc*^*GFSTF*^ is due to loss of fertility in males, females or both, we separately mated *crc* mutant females to healthy control (genotype; *yw*) males and vice versa. We observed that while *crc*^*GFSTF*^ and *crc*^*GFSTF*^/*crc*^*1*^ males produced viable progeny at similar rates to control *yw* males (data not shown), *crc* mutant females showed ∼50% reduction in egg laying in comparison to control females (Fig. 6a), again with very few of the eggs laid hatching to first instar larvae. Upon closer observation, we saw defects in the dorsal appendages of eggs laid by *crc* mutant females, with mild phenotypes such as shortening of the appendages to complete absence of one or both appendages (Fig. 6b). These data indicated that the fertility defects in *crc* mutants were due to the loss of crc function in female flies.

**Fig. 6.**
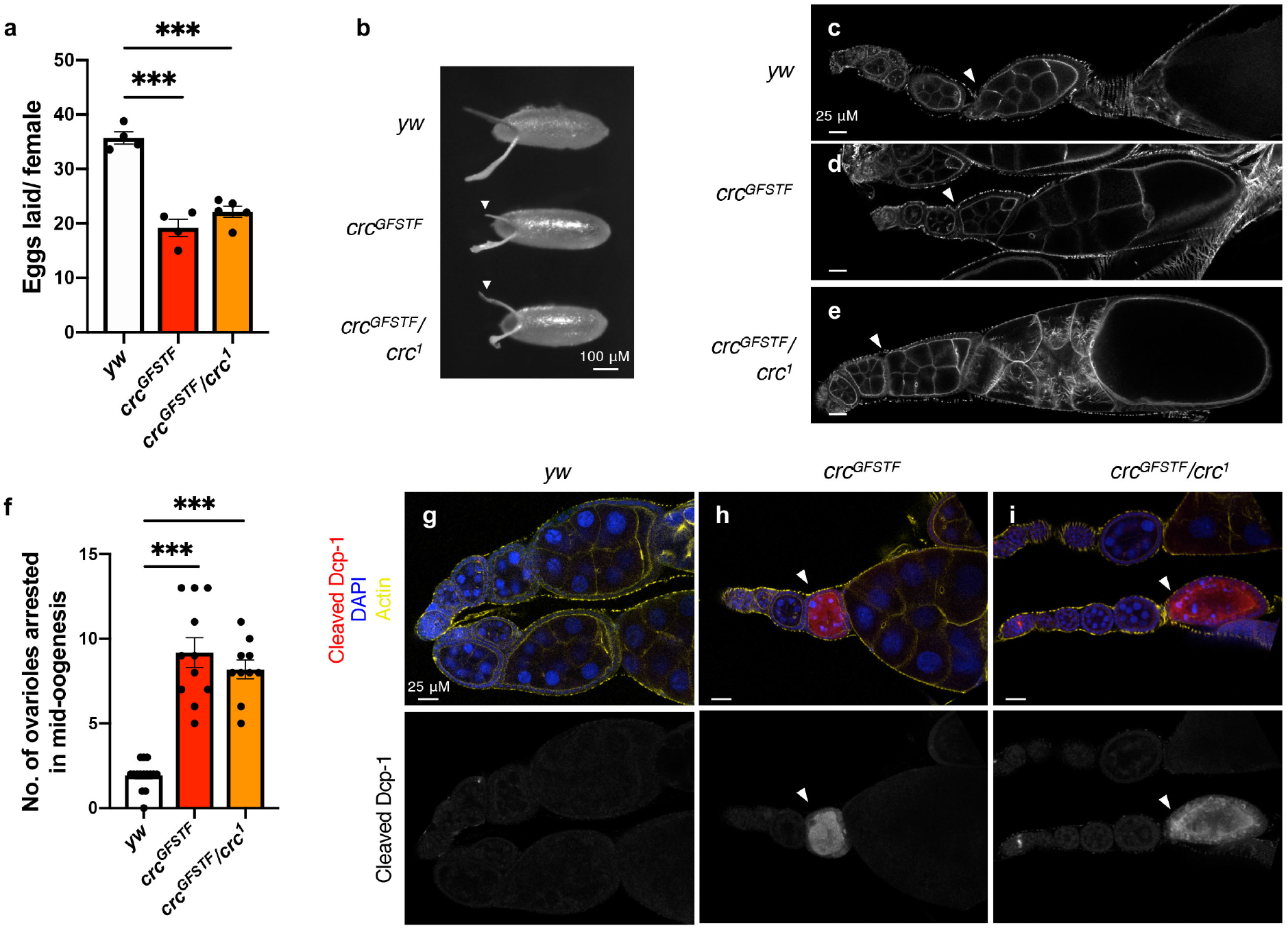
*crc* mutant females show reduced fertility due to a block in oogenesis. a. Number of eggs laid per female in a 24-hour period for control (*yw*), and *crc* mutants. The data are the mean from 4 independent experiments with five females per experiment, error bars represent standard error. ***= p<0.0001, calculated by a paired two-tailed *t*-test. b. Grayscale images of 0-24 hour eggs from females of the indicated genotypes. White arrowheads indicate dorsal appendage defects in eggs laid by *crc* mutant females, in comparison to well-formed and elongated dorsal appendages in eggs laid by control females (*yw*). c-e. Confocal images of individual ovarioles from indicated genotypes, counterstained with phalloidin (actin). Control ovarioles (*yw*) show clearly delineated individual egg chambers (white arrowheads, c) that are appropriately sized for each stage. *crc* mutant ovarioles (*crc*^*GFSTF*^, *crc*^*GFSTF*^*/crc*^*1*^) show enlarged stage 10 egg chambers, with no clear delineation between individual egg chambers (white arrowheads, d-e) indicating a block in oogenesis. f. The number of ovarioles per ovary showing enlarged stage 10 egg chambers, which are indicative of a mid-oogenesis arrest. Data represent the mean from individual ovaries of 11 animals, error bars represent standard error. ***= p<0.0001, calculated by an unpaired two-tailed *t*-test. g-i. Confocal images of ovaries from the indicated genotypes stained with the cell death marker cleaved Dcp-1 (red), nuclei counterstained with DAPI (blue) and phalloidin marking actin (yellow). White arrowheads point to egg chambers in *crc* mutant ovarioles (*crc*^*GFSTF*^, *crc*^*GFSTF*^*/crc*^*1*^) that show elevated Dcp-1 staining.

Dorsal appendages are specified and develop in the final stage of oogenesis. Each *Drosophila* ovary is comprised of 14-16 developing follicles called ovarioles, with germline stem cells residing at the anterior apex undergoing differentiation along the ovariole in individual egg chambers (Lobell et al. 2017). Each egg chamber represents a distinct stage in ovulation, with stage 14 representing a mature egg. To further dissect the dorsal appendage defects, we examined ovaries from *crc* mutant animals. We observed that ovaries from *crc*^*GFSTF*^ and *crc*^*GFSTF*^/*crc*^*1*^ were considerably swollen in comparison to control ovaries (Fig. S3a). Several ovarioles within *crc* mutant ovaries showed accumulation of stage 10 egg chambers, indicative of an arrest in oogenesis (yellow arrowheads in Fig. S3a). Indeed, examination of individual ovarioles from *crc* mutant ovaries counterstained for actin showed that loss of *crc* results in an abnormal arrangement of early stage egg chambers (Fig. 6c-e). While ovarioles from control animals showed sequentially staged and spaced egg chambers culminating in mature stage 14 eggs (Fig. 6c), ovarioles from *crc*^*GFSTF*^ and *crc*^*GFSTF*^/*crc*^*1*^ appeared to be arrested in stage 10, with improper spacing between egg chambers in earlier stages (white arrowheads, Fig. 6d, e). We quantified the number of ovarioles that displayed such arrest and found that more than half of *crc* mutant ovarioles (∼9) in each ovary showed stage 10 arrest in comparison to an average of 2-3 ovarioles arrested in ovaries from corresponding control animals (Fig. 6f).

To determine if the arrested egg chambers underwent subsequent cell death, we immunostained ovaries with an antibody that detects proteolytically activated (cleaved) caspase, Dcp-1 (Vasudevan and Ryoo 2016). We observed that stage 10 egg chambers from several *crc*^*GFSTF*^ and *crc*^*GFSTF*^/*crc*^*1*^ ovarioles showed strong cleaved Dcp-1 staining (Fig. 6g-i). These data strongly suggest that the decrease in fertility in *crc*^*GFSTF*^ and *crc*^*GFSTF*^/*crc*^*1*^ females is associated with cell death in arrested egg chambers during oogenesis.

To examine which cell types express crc in the ovary, we immunostained ovaries with GFP antibody to detect crc-GFP. However, we were unable to detect crc-GFP in this tissue (Fig. S3b, c), suggesting that crc may regulate ovulation non-autonomously. We also attempted western blotting of ovary extracts to detect crc-GFP but did not observe any detectable signal (data not shown). A previous study had suggested a non-autonomous role for fat body *Gcn2* in the regulation of oogenesis (Armstrong et al. 2014). Consistent with these observations, we were able to detect high levels of crc-GFP fusion protein in adult abdominal fat tissues from *crc*^*GFSTF*^ animals (Fig. S3d,e). These data raise the possibility that crc mediates Gcn2-signaling in fat tissues to non-autonomously regulate oogenesis.

## Discussion

ISR signaling is associated with various pathological conditions, but the role of *Drosophila crc* in adult tissues had remained unclear. This may be in part because the cytogenetic location of *crc* is very close to FRT40, and therefore, attempts to study crc function using conventional genetic mosaics have been unsuccessful. Thus far, our understanding of the role of crc in adult *Drosophila* tissues has entirely relied on RNAi experiments. Loss-of-function mutants, however, allow for unbiased discovery of developmental phenotypes, as is exemplified in our present study where we examined the role of crc in later developmental stages, adult tissues and during aging.

Generally, ER stress-imposing proteins such as Rh1^G69D^ are thought to activate the PERK-mediated ISR response amongst other ER stress responses (Donnelly et al. 2013). It is worth noting here that while both *Drosophila* and mouse models of adRP describe a protective role for *Perk* in retinal degeneration (Chiang et al. 2012; Athanasiou et al. 2017; Vasudevan et al. 2020), there has been conflicting evidence on the role of ATF4 in the mouse adRP model (Bhootada et al. 2016). In this study, we show that loss of *crc* accelerates the age-related retinal degeneration in a *Drosophila* model of adRP. As we have previously shown that *Perk* mutants similarly accelerate retinal degeneration in this model (Vasudevan et al. 2020), we interpret that *crc* mediates the effect of *Perk* in this model. Our data finds that loss of crc renders photoreceptor susceptible to retinal degeneration with age in otherwise wild type animals (solid red line, Fig. 4a). Along with our observation showing an increase in crc protein levels in older flies (Fig. 4b, c), these data indicate that photoreceptors have physiological stress that requires crc for their survival during aging.

One of the visible phenotypes in adult *crc* mutants is ectopic wing venation (Fig. 5). It has previously been demonstrated that *Gcn2* depletion in the posterior compartment of imaginal discs results in ectopic wing vein formation (Malzer et al. 2010). The study proposed that GCN2 regulates BMP signaling by modulating mRNA translation in wing discs via eIF2α phosphorylation and 4E-BP induction. Our results are consistent with this proposal since 4E-BP is a transcription target of *crc*. In addition, we report here that *crc* loss affects wing size, a finding that has not been reported previously. Given that BMP signaling has also been extensively implicated in determining wing size (Gibson and Perrimon 2005; Shen and Dahmann 2005), it is possible that GCN2-crc signaling regulates wing size via BMP signaling. It is equally possible that GCN2-crc signaling affects tissue size through its role in regulating amino acid transport and metabolism through autonomous and non-autonomous means.

*Drosophila* fat body is an organ that orchestrates organismal metabolism in response to changes in nutrient availability. While wing development is not known to be sexually dimorphic, fat tissues are known to have sex-specific effects, with particularly profound effects on female fertility is in flies and in all other sexually dimorphic organisms (Valencak et al. 2017). It has been previously demonstrated that loss of *crc* in *Drosophila* larvae leads to reduced fat content and increased starvation susceptibility (Seo et al. 2009). Correlating with this, it had been found that starvation causes effector caspase activation and cell death during mid-oogenesis (McCall 2004; Hou et al., 2008; Jenkins et al., 2013). These observations prompt us to speculate that the caspase-mediated block in oogenesis in *crc* mutants (Fig. 6) may be due to metabolic changes in the female fat body. This hypothesis integrates well with our data showing high crc activity in adult fat tissues (Fig. S3d, e) and observations from a previous study that amino acid sensing by GCN2 in *Drosophila* adult adipocytes regulates germ stem cells in the ovary (Armstrong et al. 2014). However, it remains possible that crc acts autonomously in the ovary but is undetectable using our current methods (Fig. S3b, c).

In summary, our study has found utilities for the *crc*^*GFSTF*^ allele in discovering a new role for ISR signaling in disease models and during development, and as an endogenous reporter for ISR activation.

## Methods

### Fly husbandry

Flies were reared on cornmeal-molasses media at 25°C under standard conditions except for retinal degeneration experiments when they were reared under constant light. All fly genotypes and sources used in the study are listed in Table S1.

### Phenotype analysis

Lethal phase analysis was performed as described previously (Vasudevan et al. 2020). Right wings were severed from 1-4 day old flies and imaged using a Nikon SMZ1500 microscope outfitted with a Nikon 8MP camera with NIS-Elements software. Wing size was measured using regions of interest (ROI) feature in ImageJ software.

Female fertility was quantified by placing five 1-4 day old virgin females with five *yw* males in a vial containing standard media enhanced with yeast to encourage egg laying. After allowing a day for acclimatization, the flies were moved to a new vial and the number of eggs laid in a 24-hour period were counted and quantified. Eggs were imaged for Fig. 5b by placing them on an apple juice plate and captured with the Nikon SMZ1500 microscope outfitted with 8MP Nikon camera controlled by NIS elements software. Ovaries from female flies in this experiment were dissected in cold PBS and similarly imaged on apple juice plates for Fig. S3a.

### qPCR analysis

Total RNA was prepared using TriZol (Invitrogen) from five wandering third instar larva, and cDNA was generated using random hexamers (Fisher Scientific) and Maxima H minus reverse transcriptase (Thermo Fisher) according to manufacturer’s protocol. qPCR was performed using PowerSYBR Green Mastermix (Thermo Fisher) using the following primers

crc-Fwd: GGAGTGGCTGTATGACGATAAC Rev: CATCACTAAGCAACTGGAGAGAA

Thor-Fwd: TAAGATGTCCGCTTCACCCA Rev: CGTAGATAAGTTTGGTGCCTCC

Rpl15-Fwd: AGGATGCACTTATGGCAAGC Rev: CCGCAATCCAATACGAGTTC

### Immunostaining

Ovaries and fat bodies were dissected in cold PBS from female flies reared for 2-3 days along with *yw* males on standard media enhanced with yeast. Tissues were fixed in 4% PFA in PBT (0.2% Triton-X 100, 1X PBS) for 30 minutes, washed 3x with PBT, and blocked in 1% BSA, PBT for 3 hours, all at room temperature. Tissues were stained overnight at 4°C with the primary antibodies diluted in PBT, following which they were washed 3X with PBT and incubated with AlexaFluor-conjugated secondary antibodies (Invitrogen) in PBT for 3 hours at room temperature. Tissues were mounted in 50% glycerol containing DAPI.

Eye imaginal discs were dissected from wandering 3^rd^ instar larva in cold PBS and fixed in 4% PFA in PBS for 20 minutes, following which they were washed 2x with PBS and permeabilized in 1X PBT for 20 minutes, all at room temperature. Discs were incubated in primary antibodies diluted in PBT for 2 hours, washed 3x in PBT, incubated in AlexaFluor-conjugated secondary antibodies (Invitrogen) in PBT for 1 hour and washed 3x in PBT prior to mounting in 50% glycerol containing DAPI.

#### Antibodies

Phalloidin-Alexa647 (1:1000, Invitrogen), chicken anti-GFP (1:500, Aves Labs), Rabbit anti-cleaved Dcp-1 (1:100, Cell Signaling), Mouse anti-4C5 for Rh1 (1:500, DSHB), Rabbit anti-eIF2α (1:500, AbCam), rabbit anti-S51 peIF2α (1:500, AbCam). All images were obtained on a Zeiss LSM 700 confocal microscope with ZEN elements software and a 20X air or 40X water lens.

### Retinal degeneration

All experiments were performed in a *white* mutant background since *crc*^*GFSTF*^, *crc*^*1*^, and *ninaE*^*G69D*^, do not have eye color. 0-3 day old male flies were placed (20 animals/vial) under 1000-lumen light intensity, and their pseudopupil structures monitored under blue light at 3-day intervals for a 30-day period. Media was replaced every 3 days, and flies with disrupted pseudopupils in one or both eyes were marked as having retinal degeneration.

### Western blotting

Fly head extracts were prepared from 6 severed male fly heads in 30 μl lysis buffer containing 10mM Tris HCl (pH 7.5), 150mM NaCl, protease inhibitor cocktail (Roche), 1mM EDTA, 1% SDS. Following SDS-PAGE and western blotting, proteins were detected using primary antibodies and IRDye-conjugated secondary antibodies (LI-COR) on the Odyssey system. Primary Rabbit anti-GFP (1:500, Invitrogen) and mouse anti-Tub (1:1000, DHSB).

### Amino acid deprivation

0-3 day old female flies were placed (10 animals/vial) in standard media or in vials containing 5% sucrose, 2% agarose prepared in dH_2_O. The number of survivors was counted every 24 hours and survivors were moved to new media.

## Acknowledgements

We thank Hugo Bellen’s laboratory for making available the *crc* MIMIC RMCE line, and Drs. Lacy Barton and Lydia Grmai for discussions on the ovary phenotypes, and Drs. Erika Bach and Jessica Treisman and their laboratories for helpful discussions that improved this work. We thank the Bloomington Drosophila Stock Center (NIH P40OD018537) for supplying many of the fly stocks, and FlyBase (U41 HG000739) for curating sequence data used in this study.

## Contributions

D.V. and H.D.R. conceptualized the project, analyzed the data, and wrote the manuscript. H.K. performed all the wing phenotype analyses, G.T. executed all western blotting experiments, and D.V. performed all other experiments.

